# A SAC phosphoinositide phosphatase controls rice development via hydrolyzing phosphatidylinositol 4-phosphate and phosphatidylinositol 4,5-bisphosphate

**DOI:** 10.1101/740001

**Authors:** Tao Guo, Hua-Chang Chen, Zi-Qi Lu, Min Diao, Ke Chen, Nai-Qian Dong, Jun-Xiang Shan, Wang-Wei Ye, Shanjin Huang, Hong-Xuan Lin

**Affiliations:** National Key Laboratory of Plant Molecular Genetics, CAS Centre for Excellence in Molecular Plant Sciences and Collaborative Innovation Center of Genetics & Development, Shanghai Institute of Plant Physiology & Ecology, Shanghai Institute for Biological Sciences, Chinese Academic of Sciences, Shanghai 200032, China; University of the Chinese Academy of Sciences, Beijing 100049, China; School of Life Science and Technology, ShanghaiTech University, Shanghai, 201210, China; Center for Plant Biology, School of Life Sciences, Tsinghua University, Beijing, China

**Keywords:** Rice, SAC phosphatase, phosphatidylinositol 4-phosphate, phosphatidylinositol 4;5-bisphosphate, cell elongation

## Abstract

Phosphoinositides (PIs) as regulatory membrane lipids play essential roles in multiple cellular processes. Although the exact molecular targets of PIs-dependent modulation remain largely elusive, the effects of disturbed PIs metabolism could be employed to propose regulatory modules associated with particular downstream targets of PIs. Here, we identified the role of *GRAIN NUMBER AND PLANT HEIGHT 1* (*GH1*), which encodes a suppressor of actin (SAC) domain-containing phosphatase with unknown function in rice. Endoplasmic reticulum-localized GH1 specifically dephosphorylated and hydrolyzed phosphatidylinositol 4-phosphate (PI4P) and phosphatidylinositol 4,5-bisphosphate [PI(4,5)P_2_]. Inactivation of GH1 resulted in massive accumulation of both PI4P and PI(4,5)P_2_, while excessive GH1 caused their depletion. Notably, superabundant PI4P and PI(4,5)P_2_ could both disrupt actin cytoskeleton organization and suppress cell elongation. Interestingly, both PI4P and PI(4,5)P_2_ inhibited actin-related proteins 2 and 3 (Arp2/3) complex-nucleated actin branching networks *in vitro*, whereas PI(4,5)P_2_ showed more dramatic effect in a dose-dependent manner. Overall, the overaccumulation of PI(4,5)P_2_ resulted from dysfunction of SAC phosphatase possibly perturbs Arp2/3 complex-mediated actin polymerization, thereby disordering the cell development. These findings imply that Arp2/3 complex might be the potential molecular target of PI(4,5)P_2_-dependent modulation in eukaryotes, thereby providing new insights into the relationship between PIs homeostasis and plants growth and development.

---

Phosphoinositides (PIs) as a minor component of cell membranes and phosphorylated derivatives of phosphatidylinositol are known to liberate the second messengers, inositol 1,4,5-triphosphate and diacylglycerol in response to activation of cell surface receptors (Boss and Im, 2012). Nevertheless, emerging evidence suggests that PIs also play crucial roles as signaling molecules, functioning as regulatory lipids in various fundamental cellular processes (Heilmann, 2016). A new detection approach further confirmed that phosphatidylinositol 4-phosphate (PI4P) and phosphatidylinositol 4,5-bisphosphate [PI(4,5)P_2_] are the most abundant PIs in the cells (Balla, 2013; Kielkowska et al., 2014). PI4P is phosphorylated by phosphatidylinositol phosphate kinases, resulting in the production of PI(4,5)P_2_, while conversely PI(4,5)P_2_ is dephosphorylated by PI phosphatases to generate PI4P, supporting the hypothesis that turnover and dynamic formation of PIs is necessary for plant development (Gerth et al., 2017).

The plant actin cytoskeleton plays essential roles in cell morphogenesis, and the dynamics of actin cytoskeleton drives the membrane deformation and trafficking, which is required for polar cell growth and movement of vesicles and organelles (Pleskot et al., 2014; Wang et al., 2017). This dynamic remodeling is controlled by diverse actin-binding proteins (ABPs), which underpin various signaling pathways. Membrane PIs have also been shown to affect the dynamic assembly and disassembly of actin filaments (Li et al., 2015; Gerth et al., 2017), notably PI(4,5)P_2_ regulates the activity and distribution of ABPs, which in turn are tightly linked to actin polymerization activity (Saarikangas et al., 2010; Zhang et al., 2012; Bothe et al., 2014; Pleskot et al., 2014).

Moreover, the highly conserved Arp2/3 complex was found to nucleate branched actin filament networks, although it is dependent on stimulation of nucleation promoting factors (NPFs) such as Wiskott-Aldrich syndrome protein (WASP), and WASP family verprolin homologous-protein (WAVE)/suppressor of cAMP receptor (SCAR) (Pollard, 2007; Rotty et al., 2013). NPFs tend to share a typical carboxy-terminal tripartite functional VCA (verprolin homology, central hydrophobic, and acidic C-terminal) domain, which require PI(4,5)P_2_ to overcome autoinhibition and bind to the Arp2/3 complex for conformation activation (Pollard, 2007; Padrick and Rosen, 2010). In plants, activation of the Arp2/3 complex is relatively simple, and appears dependent on the WAVE/SCAR family and heteromeric WAVE/SCAR regulatory complex as a sole trigger (Frank et al., 2004; Yanagisawa et al., 2013). Nevertheless, precise cellular control of WAVE/SCAR-Arp2/3 module-nucleated actin polymerization and the relationship with membrane PIs remains unknown.

In general, the reversible phosphorylation of PIs dominated by a set of kinases and phosphatases controls the turnover and dynamics of membrane PIs (Hsu and Mao, 2013; Takasuga and Sasaki, 2013). PI phosphatases such as inositol polyphosphate phosphatases and SAC domain containing-phosphatases were also found to hydrolyze phosphate from PIs, thereby modifying PI levels (Zhong and Ye, 2003; Ercetin and Gillaspy, 2004; Zhong et al., 2005; Thole et al., 2008; Novakova et al., 2014; Gerth et al., 2017). However, the cellular functions of the SAC phosphatases involved in PI metabolism during plant growth and development remain largely unknown in crops. This study carried out functional characterization of an unknown SAC phosphatase in rice that is required for homeostasis of PI(4,5)P_2_ and PI4P and thus affects plant cell morphogenesis. These results implicate that overaccumulation of PI(4,5)P_2_ possibly disrupts the Arp2/3 complex activity and actin cytoskeleton organization, thereby suppressing cell elongation in rice.

## Results

### *GH1* contributes to rice plant and panicle development

To determine the molecular basis of panicle morphogenesis in rice, parental varieties Guichao-2 (GC) and CB, which exhibit obvious differences in plant and panicle architecture, were selected to identify new quantitative trait loci (QTLs) (Supplemental Fig. S1, A-H). GC and CB were crossed to produce an F_2_ mapping population and then a map-based cloning method was used to isolate a new gene locus for grain number per panicle located in the region of chromosome 2 defined using molecular markers NB-2-80 and NB-2-93 (Fig. 1A). These findings coincide with the results of bulked segregation analysis-assisted mapping (Supplemental Fig. S2) (Takagi et al., 2013). Therefore, suggesting that a single locus, referred to here as *GRAIN NUMBER AND PLANT HEIGHT 1* (*GH1*), is pleiotropically responsible for plant and panicle morphogenesis. The *GH1* gene was then fine mapped to a region between marker loci NB-2-83c and NB-2-83d using a larger segregating F_2_ population (Fig. 1A). This region contained four predicted genes, but sequence comparisons using the region from the mapping parents showed that only Os02g0554300 was polymorphic. It was therefore selected as the candidate gene of *GH1*, in which a C to T nucleotide mutation from the CB genotype resulted in premature stop of this protein (Fig. 1A).

**Figure 1.**
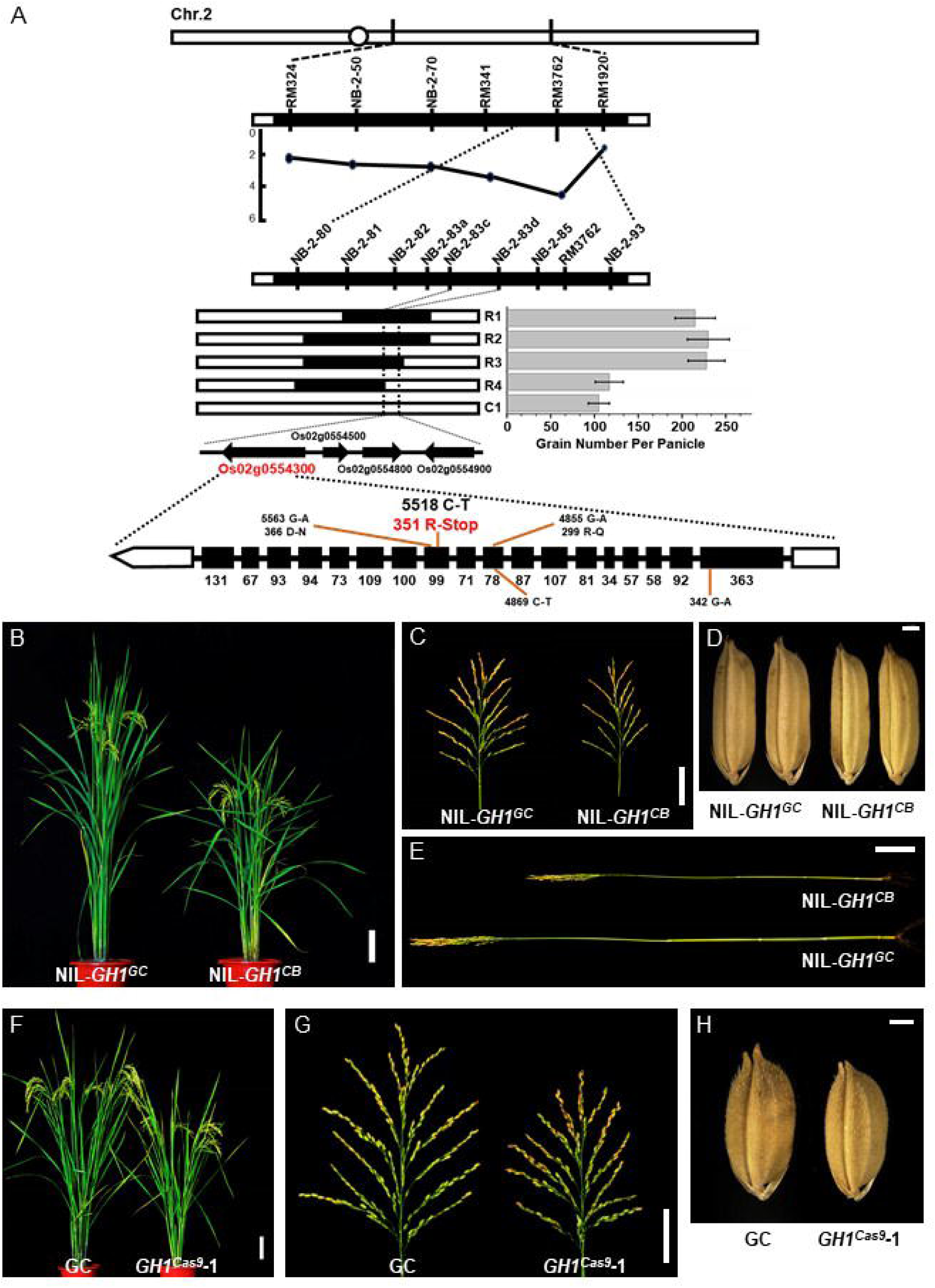
*GH1* contributes to rice plant and panicle development. A, Map-based cloning of *GH1*. Grain number per panicle of the three recombinant lines (R1-R3) was higher than that of recombinant line R4 and the control (C1, homozygous for CB in the target region) (n=15 plants, values represent the mean ± SD). Filled and open bars represent homozygous chromosomal segments for GC and CB, respectively. Four open reading frames were found in the candidate region, Os02g0554300, Os02g0554500, Os02g0554800 and Os02g0554900. Sequencing of the full-length of the four genes revealed a C-T mutation in the 11^th^ exon of Os02g0554300, resulting in premature stop of this protein. B, Plant architecture of NIL-*GH1^GC^* and NIL-*GH1^CB^* at the reproductive phase. Scale bar, 10 cm. C, Mature panicles of NIL-*GH1^GC^* and NIL-*GH1^CB^*. Scale bar, 5 cm. D, Mature grains from NIL-*GH1^GC^* and NIL-*GH1^CB^*. Scale bar, 1 mm. E, The culms of NIL-*GH1^GC^* and NIL-*GH1^CB^*. Scale bar, 5 cm. F, Plant architecture of GC and *GH1^cas9^-*1 at the reproductive phase. Scale bar, 10 cm. G, Mature panicles of GC and *GH1^cas9^-*1. Scale bar, 5 cm. H, Mature grains of GC and *GH1^cas9^-*1. Scale bar, 1 mm.

To estimate the effects of *GH1*, a nearly isogenic line (NIL) containing a 50-kb chromosomal region of GC at the *GH1* locus was developed under the CB genetic background. NIL-*GH1^CB^* displayed a distinct short stem length and reduced panicle and grain size compared with NIL-*GH1^GC^* (Fig. 1, B-E). Accordingly, the average plant height, grain number per panicle, grain length and width of NIL-*GH1^CB^* decreased significantly (Supplemental Fig. S1, I-L). The identity of *GH1* was then validated by editing the candidate gene using the CRISPR/Cas9 method (Ma et al., 2015), revealing a similar phenotype between *GH1* knockout line *GH1^cas9^* and NIL-*GH1^CB^*, which had smaller panicles and grains and was shorter in height (Fig. 1, F-H; Supplemental Fig. S3, A-G and S3, J-M). Overexpression of *GH1* also resulted in weaker growth of plant and panicle architecture compared with the wild-type (Supplemental Fig. S4, A-H). Overall, these results suggest that *GH1* plays a crucial role in plant and panicle development during rice morphogenesis.

### GH1 phosphatase specifically dephosphorylates PI4P and PI(4,5)P_2_

*GH1* was therefore thought to encode a previously unknown SAC domain-containing phosphatase in rice that is conserved throughout the plant kingdom (Supplemental Fig. S5). Notably, mutation of arginine at position 351 directly resulted in premature stop of GH1^CB^ at the CB allele and deletion of the core catalytic motif in SAC phosphatase (Fig. 2A). Recently, SAC domain-containing proteins were found to possess PI phosphatase activity in eukaryotic species (Wei et al., 2003; Liu et al., 2008; Thole et al., 2008; Brice et al., 2009; Lee et al., 2011; Hsu and Mao, 2013). Although the SAC domain is homologous among different proteins, they appear to display varied substrate specificity and subcellular localization (Hsu and Mao, 2013). Nevertheless, the specific functions of SAC phosphatase in crops remain largely unknown. Thus, to elucidate the phosphatase activity and substrate specificity of GH1, GH1^GC^ and GH1^CB^ fusion proteins from Sf9 insect cells were purified for *in vitro* phosphatase activity assay. Malachite green phosphate assay was then used to determine the enzyme activity of PI phosphatase. Interestingly, GH1^GC^ protein exhibited preference and substrate specificity towards PI4P and PI(4,5)P_2_ rather than PI3P, PI5P, and PI(3,4)P_2_ compared with the control, with a higher level of PI4P phosphatase activity *in vitro* (Fig. 2B). Moreover, GH1^CB^ failed to recognize the diverse PI isoforms, especially PI4P and PI(4,5)P_2_, suggesting that mutation of GH1^CB^ disabled its phosphatase activity (Fig. 2B). Furthermore, to test whether GH1 binds to phospholipids directly, the fusion proteins were purified and subjected to a protein-lipid overlay assay on which 15 different lipids were spotted (Fig. 2C). The results showed that the fusion protein GH1^GC^ can strongly bind to PI4P and PI(4,5)P_2_, and show weak binding to PI(3,4,5)P_3_; however, the mutated GH1^CB^ hardly binds to the phospholipids (Fig. 2D). These results also coincide with the previous suggestion that the conserved catalytic site of the SAC domain consisting of a peptide motif with a CX_5_R sequence at the C-terminal plays an essential role in phosphate hydrolysis in yeast (Manford et al., 2010). Overall, these findings suggest that the SAC phosphatase GH1 specifically dephosphorylates and hydrolyzes PI4P and PI(4,5)P_2_.

**Figure 2.**
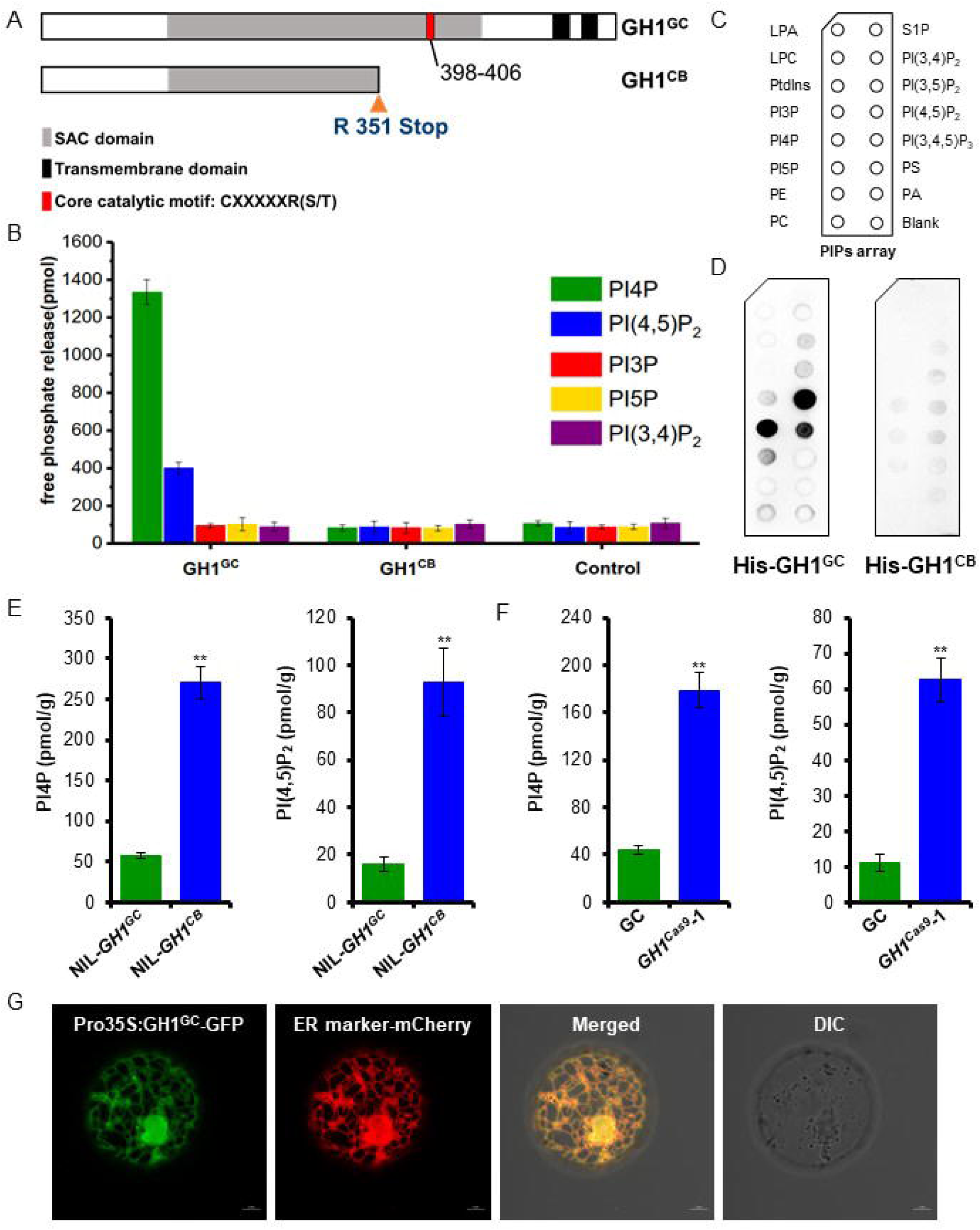
GH1 phosphatase specifically dephosphorylates PI4P and PI(4,5)P_2_. A, C-T mutation of the *GH1* allele from CB caused a premature stop codon that resulted in absence of the core catalytic motif and two transmembrane domains in the C-terminal of the predicted SAC domain-containing phosphatase. B, The SAC domain-containing GH1^GC^ phosphatase showed high substrate specificity towards PI4P and PI(4,5)P_2_ over other PIs isoforms, while the mutated GH1^CB^ showed no phosphatase activity as a control (n=3 biologically independent reactions). Values represent the mean ± SD. C, Schematic diagram of a PIP strip containing an array of immobilized phospholipids: lysophosphatidic acid (LPA), lysophosphocholine (LPC), phosphatidylinositol (PtdIns), PI3P, PI4P, PI5P, phosphatidylethanolamine (PE), phosphatidylcholine (PC), sphingosine-1-phosphate (S1P), PI(3,4)P_2_, PI(3,5)P_2_, PI(4,5)P_2_, PI(3,4,5)P_3_, phosphatidylserine (PS), and phosphatidic acid (PA). D, Purified recombinant His-GH1^GC^ (left) and His-GH1^CB^ (right) were overlaid onto PIP strip membranes. Proteins bound to lipids were detected by immunoblotting with anti-His monoclonal antibody. E and F, Comparisons of endogenous levels of PI4P and PI(4,5)P_2_ extracted from rice leaf blades of NIL-*GH1^GC^* and NIL-*GH1^CB^* (E) and GC and *GH1^cas9^-*1 plants (F) (n=3 biologically independent assays). Values represent the mean ± SD. \*\**P* < 0.01 compared with the control using the Student’s *t*-test. G, Subcellular localization of GH1 in rice protoplasts. The GH1 protein was shown to target the ER by transient expression of GH1^GC^-GFP, which merged with the ER marker-mCherry. DIC, differential interference contrast. Scale bar, 5 μm.

To further investigate the effect of GH1 on PI4P and PI(4,5)P_2_ metabolism in rice, endogenous levels of PI4P and PI(4,5)P_2_ were determined using enzyme-linked immune sorbent assay (ELISA) with different plant leaves. Both PI4P and PI(4,5)P_2_ levels were markedly higher in NIL-*GH1^CB^* than NIL-*GH1^GC^* (Fig. 2E), suggesting that loss-of-function of *GH1* results in massive accumulation of these PI isoforms in rice. Consistent with this, *GH1* knockout line *GH1^cas9^* also showed a significant increase in endogenous PI4P and PI(4,5)P_2_ compared with the wild-type (Fig. 2F; Supplemental Fig. S3, H and I). In contrast, *GH1^OE^* overexpressing plants had reduced PI4P and PI(4,5)P_2_ levels (Supplemental Fig. S4, I and J). These findings suggest that GH1 specifically acts as a PI4P and PI(4,5)P_2_ phosphatase, maintaining degradation and homeostasis of PI4P and PI(4,5)P_2_, and thereby acting as an essential regulator of membrane PI signaling-networks in rice. Membrane lipids can be transferred between bilayers at contact sites between the endoplasmic reticulum (ER) and other membranes to maintain homeostasis (Stefan et al., 2013; Holthuis and Menon, 2014; Chung et al., 2015). In previous reports, counter transport and exchange of PI4P and phosphatidylserine (PS) between the ER and plasma membrane (PM) enabled delivery of PI4P to the ER during degradation and synthetic PS from the ER to the PM, controlling PI4P levels and selectively enriching PS in the PM (Stefan et al., 2011; Chung et al., 2015). In both yeast and mammalian cells, the PI4P phosphatase Sac1 is localized in the ER and Golgi apparatus where it acts as an integral membrane protein, and PI4P is hydrolyzed by the ER protein Sac1 (Stefan et al., 2011; Mesmin et al., 2013). These finding suggest that GH1 also might function as an ER protein in rice, with two tandem transmembrane domains in its C-terminal (Supplemental Fig. S6). Subcellular localization of GH1 in rice protoplasts was therefore examined using fluorescence signals from green fluorescent protein (GH1-GFP) fusion protein. Signals coincided with those of the ER-marker protein (Nelson et al., 2007), suggesting that the SAC phosphatase GH1 degrades PI4P and PI(4,5)P_2_ within the ER (Fig. 2G).

### *GH1* is required for actin cytoskeleton organization and organelle development in rice

To understand the cellular roles of the SAC phosphatase GH1, vertical sections of the central part of the culm were therefore compared between NIL-*GH1^GC^* and NIL-*GH1^CB^* using X-ray microscopy to determine the cellular basis for the short stem. Observations revealed that the average cell length of the culm was significantly smaller in NIL-*GH1^CB^* than NIL-*GH1^GC^*, whereas the cell width was comparable (Fig. 3, A and B). These findings suggest that *GH1* contributes to cell morphogenesis in rice, with deletion suppressing cell elongation. It has been suggested that SAC domain-containing proteins influence organization of the actin cytoskeleton in yeast and plant cells (Foti et al., 2001; Zhong et al., 2005). To explore whether *GH1* is also associated with actin cytoskeleton modulation in rice, the organization of actin filaments in the root tip cells was also examined. Root tip cells from parent GC and NIL-*GH1^GC^* plants formed a structured actin filaments cable network, while in CB and NIL-*GH1^CB^*, the actin filaments appeared to have lost their organized structure (Fig. 3C), suggesting that *GH1* is responsible for fine organization of the actin cytoskeleton. Actin filament orientation in all directions (360°) was therefore examined using visual statistical analysis, revealing a specific direction in GC and NIL-*GH1^GC^*. Meanwhile, in CB and NIL-*GH1^CB^*, the actin cytoskeleton displayed disordered distribution without definite orientation (Fig. 3D). These findings imply that homeostasis of membrane PI4P and PI(4,5)P_2_ under GH1 control could be required for actin cytoskeleton organization in rice.

**Figure 3.**
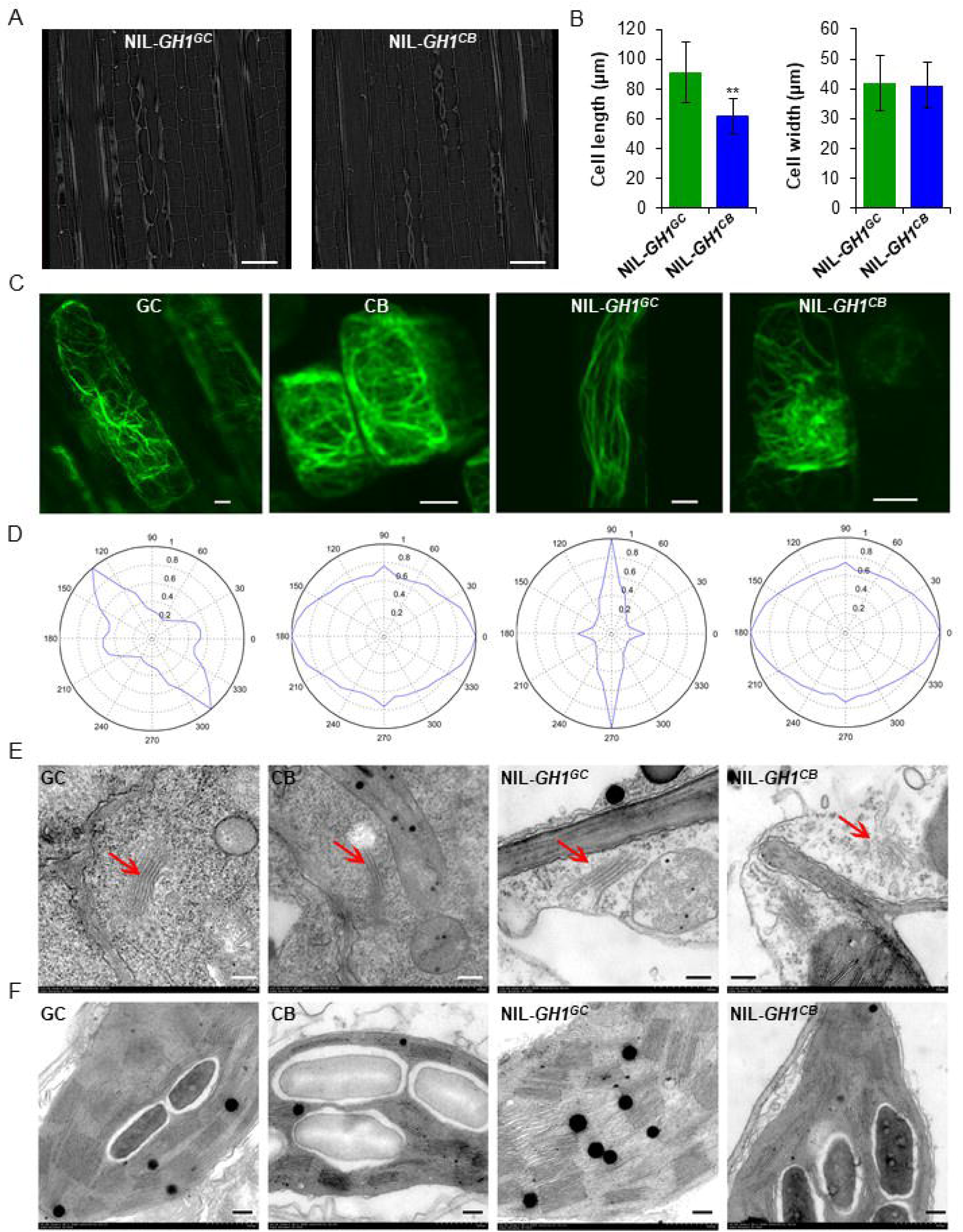
*GH1* is required for actin cytoskeleton organization and organelle development in rice. A, Scanning of vertical sections of the rice culm from NIL-*GH1^GC^* and NIL-*GH1^CB^* using Xradia 510 versa 3D X-ray microscopy. Scale bar, 100 μm. B, Comparisons of cell length and width within the culm of NIL-*GH1^GC^* and NIL-*GH1^CB^* (n=10 views). Values represent the mean ± SD. \*\**P* < 0.01 compared with NIL-*GH1^GC^* using the Student’s *t*-test. C, F-actin observations of rice root tip cells from GC, CB, NIL-*GH1^GC^*, and NIL-*GH1^CB^* stained with ActinGreen 488. Scale bar, 5 μm. D, Visual statistical analysis of actin filament orientation and actin cytoskeleton arrangement in all directions (360°) using Discrete Fourier Transform. E and F, Observations of Golgi apparatus (E) and chloroplasts (F) from GC, CB, NIL-*GH1^GC^*, and NIL-*GH1^CB^* using transmission electron microscopy. Scale bar, 2 μm.

A previous study revealed that lipid phosphatase, in regulating PI homeostasis, plays an essential role in Golgi membrane organization in mammals (Liu et al., 2008). However, whether SAC phosphatases play a similar biological role in plants remains unknown. Golgi morphology in rice leaf cells was therefore examined by transmission electron microscopy. Intact multilayered Golgi apparatus were observed in the GC and NIL-*GH1^GC^* cells; however, in CB and NIL-*GH1^CB^*, they were impaired with concomitant collapse and a vesicle-like structures (Fig. 3E). These findings therefore suggest a potential role of *GH1* in Golgi development and vesicle trafficking in plants. Moreover, they also suggest that elevated PI4P and PI(4,5)P_2_ levels could disrupt the structural integrity and function of Golgi apparatus.

Chloroplast morphology in the leaf cells was subsequently examined, revealing a dramatic reduction in thylakoid layers and the number of grana in CB and NIL-*GH1^CB^* compared with GC and NIL-*GH1^GC^* (Fig. 3F). These findings suggest that increased levels of membrane PI4P and PI(4,5)P_2_ could also disrupt thylakoid formation and grana assembly during chloroplast development in rice. Interestingly, this result coincides with reported evidence whereby inhibition of PI4P synthesis accelerates chloroplast division, supporting the idea that PI4P negatively regulates chloroplast division in *Arabidopsis* (Okazaki et al., 2015). Taken together, these findings suggest that the massive accumulation of membrane PI4P and PI(4,5)P_2_ resulting from terminated GH1 production could disturb organelle development and cytoarchitecture.

### PI(4,5)P_2_ specifically inhibits Arp2/3 complex-mediated actin polymerization *in vitro*

Overall, the present findings imply that maintaining proper balance between PI4P and PI(4,5)P_2_ levels via GH1 phosphatase is critical to cytoskeletal organization and organelle integrity. It was therefore hypothesized that the collapsed organelles resulting from overaccumulation of PI4P and PI(4,5)P_2_ were associated with disruption of the actin cytoskeleton. Nevertheless, the molecular mechanisms underlying PI4P and PI(4,5)P_2_-induced modulation of the actin cytoskeleton remain elusive. Surprisingly, PI4P and PI(4,5)P_2_ were observed to be involved in the Arp2/3 complex-mediated actin polymerization *in vitro* (Supplemental Fig. S7). The highly conserved Arp2/3 complex functions as a nucleator of actin filament dynamics in a wide range of eukaryotic cells, which consists of seven subunits: two actin-related proteins, Arp2 and Arp3, stabilized in an inactive state by five other subunits (Pollard, 2007). However, the precise function of Arp2/3 complex in plant cells remains enigmatic, and especially little is known about the counterparts and distinction of Arp2/3 complex in rice. To determine whether PI4P and PI(4,5)P_2_ are crucial lipid molecules for Arp2/3 complex action, Arp2/3 activity was therefore assayed by spectrofluorimetry and total internal reflection fluorescence microscopy (TIRFM) with the purified porcine Arp2/3 complex which is accepted and generally used for eukaryotic assay. The Arp2/3 complex stimulated actin polymerization 10-fold in conjunction with the indispensable VCA domain of WASP (Fig. 4A). However, both PI4P and PI(4,5)P_2_ inhibited Arp2/3 complex-mediated actin polymerization (Supplemental Fig. S7A), in which PI(4,5)P_2_ had more dramatic effect in a dose-dependent manner despite the analogous structures of PI(4,5)P_2_ and PI4P (Fig. 4A; Supplemental Fig. S7, A and B). These findings suggest that PI(4,5)P_2_ could inhibit the VCA-induced Arp2/3 complex activity. As expected, TIRFM further revealed that activation of the Arp2/3 complex via the VCA protein resulted in the generation of branched actin filaments, a hallmark of Arp2/3 complex-mediated actin filament nucleation (Fig. 4B). However, fewer actin branched junctions formed in the presence of both PI(4,5)P_2_ and PI4P, most notably PI(4,5)P_2_ (Fig. 4B; Supplemental Video S1), suggesting that PI(4,5)P_2_ could specifically inhibit Arp2/3 complex-mediated actin polymerization. Moreover, the elevated VCA proteins partly released the PI(4,5)P_2_-inhibited kinetics of actin polymerization (Supplemental Fig. S7C). Accordingly, a pull-down assay showed that PI(4,5)P_2_ could compete with VCA proteins binding to the Arp2/3 complex (Fig. 4C), suggesting that PI(4,5)P_2_ maybe acts as a competitive inhibitor of the NPFs-activated Arp2/3 complex. Overall, these findings imply that although PI(4,5)P_2_ was only observed *in vitro* to inhibit Arp2/3 complex-mediated actin polymerization, the Arp2/3 complex might function as the molecular target of PI(4,5)P_2_-dependent regulation.

**Figure 4.**
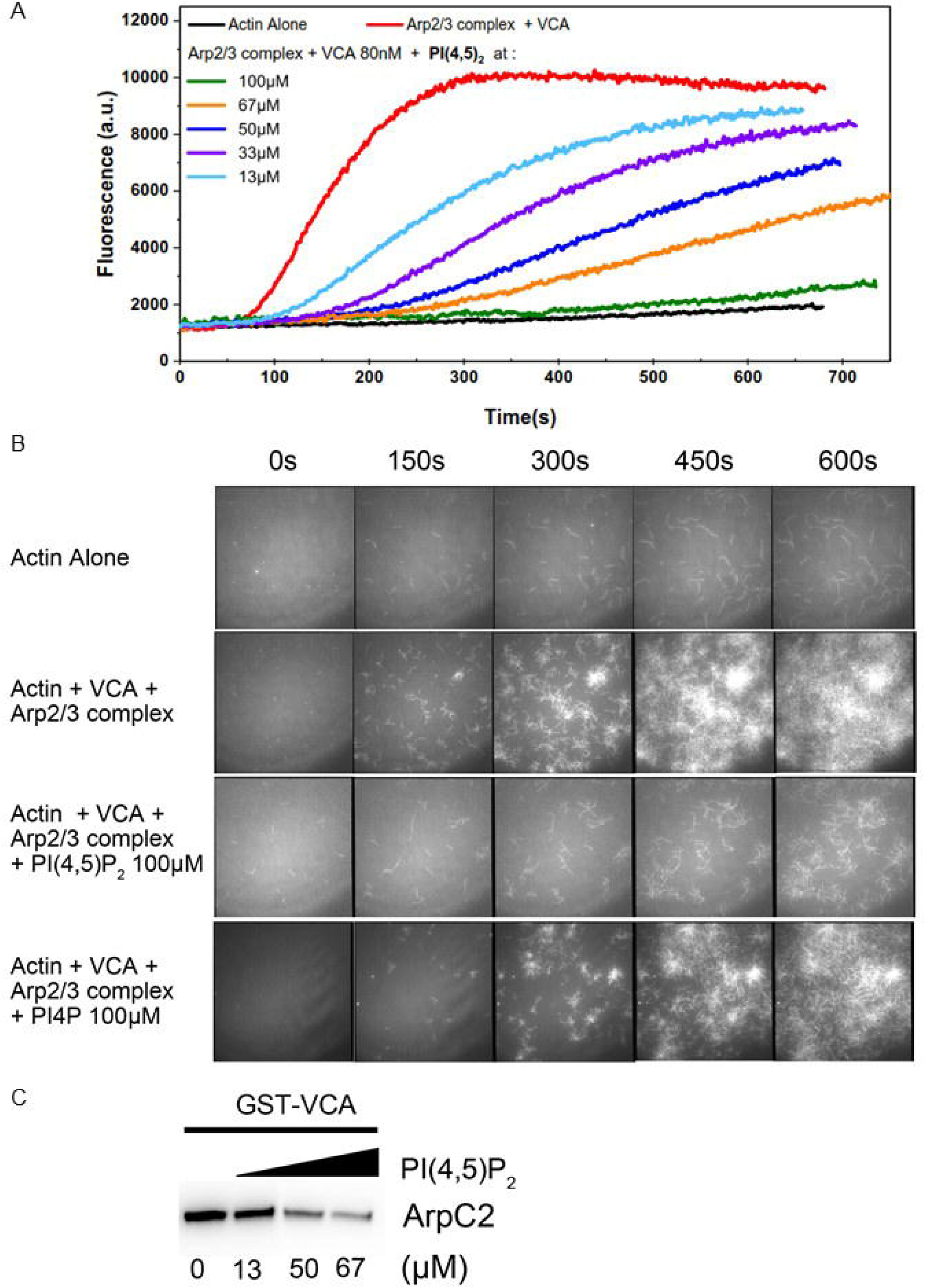
PI(4,5)P_2_ specifically inhibits Arp2/3 complex-mediated actin polymerization *in vitro*. A, Spectrofluorimetry assay using pyrene-labelled actin to monitor polymerization. PI(4,5)P_2_ inhibited VCA-promoted Arp2/3 complex activity in a dose-dependent manner. a.u., arbitrary units. B, Actin polymerization monitored by TIRFM. In the presence of PI(4,5)P_2_, the branches initiated by the Arp2/3 complex were dramatically inhibited, whereas its structural analogue, PI4P, had a much weaker inhibiting effect. C, Pull-down assay of the Arp2/3 complex with GST-VCA under elevated concentrations of PI(4,5)P_2_. Anti-ArpC2 was used for the immunoblotting assay. ArpC2 is a subunit of the Arp2/3 complex.

## Discussion

Based on the present results, it was hypothesized that massive accumulation of PI(4,5)P_2_ could disrupt organization of Arp2/3 complex-mediated actin polymerization, suppressing cell elongation. A working model was subsequently proposed to describe the role of GH1 phosphatase-controlled membrane PI(4,5)P_2_ in cell morphogenesis. Briefly, ER-localized GH1^GC^ could specifically hydrolyze membrane PI4P and PI(4,5)P_2_, maintaining PI dynamics and PI(4,5)P_2_ distribution on the cell PM. Generally, the normal activated Arp2/3 complex initiates actin polymerization at the cell cortex under relatively low PI(4,5)P_2_ levels, nucleating the branched actin filament network for specific formation of the actin cytoskeleton (Fig. 5, A and B). However, the GH1^CB^ mutation promotes endogenous PI(4,5)P_2_ levels in the PM, thus inhibiting Arp2/3 complex activity and disrupting actin polymerization. Failed nucleation of the branched actin network at the cell cortex subsequently might disrupt organization of the actin cytoskeleton in the plant cell, leading to dysfunctional Golgi apparatus and chloroplasts, in turn suppressing plant cell development (Fig. 5, A and B).

**Figure 5.**
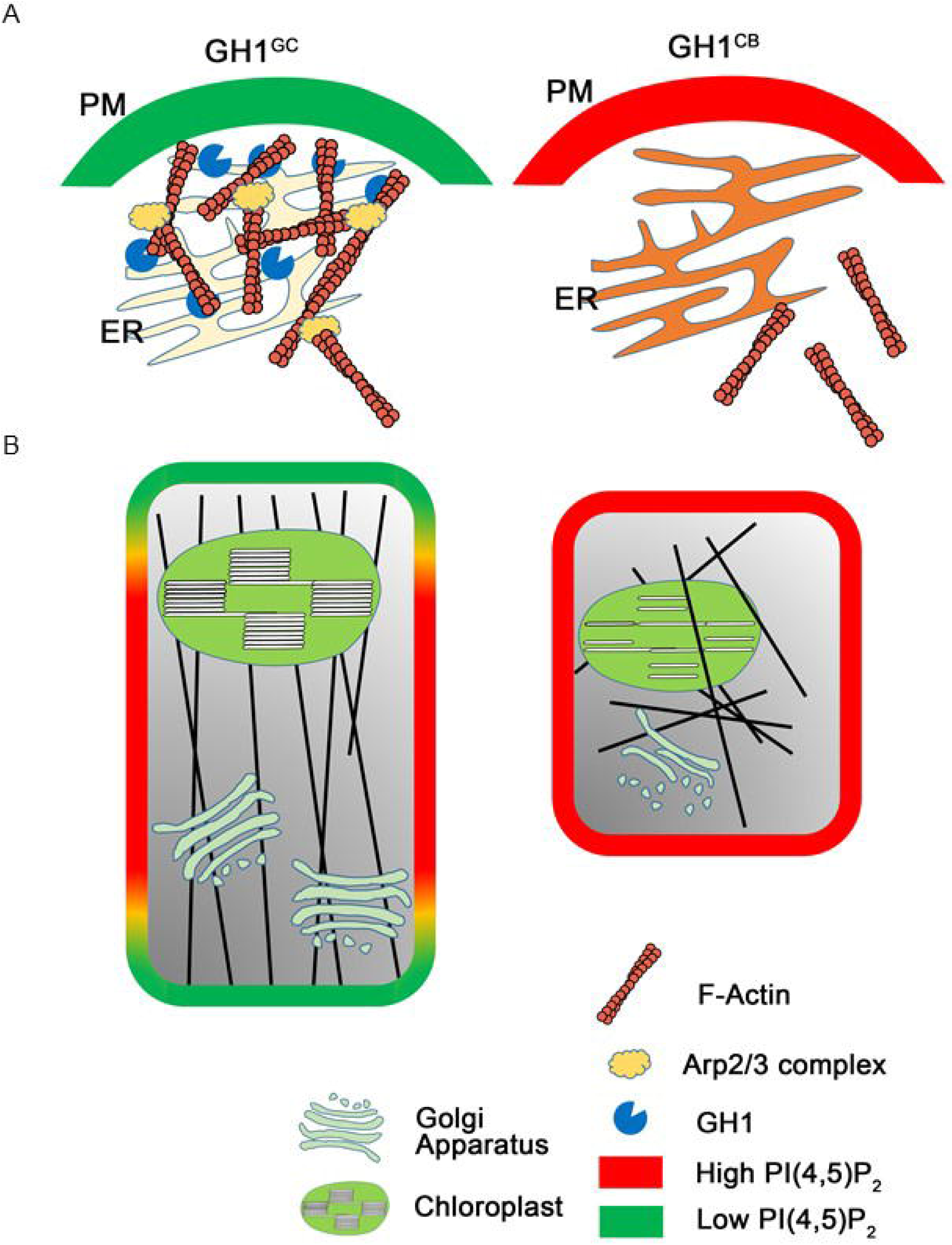
A proposed working model of the role of GH1 phosphatase-controlled membrane PI(4,5)P_2_ in actin cytoskeleton organization during cell morphogenesis. A, ER-localized GH1^GC^ specifically hydrolyzes membrane PI4P and PI(4,5)P_2_, maintaining PI dynamics, including levels of PI(4,5)P_2_ in the plasma membrane of plant cells. The activated Arp2/3 complex then could initiate actin polymerization in the cell cortex at relatively low PI(4,5)P_2_ levels. However, mutation of GH1^CB^ promoted endogenous PI(4,5)P_2_ levels at the plasma membrane, directly inhibiting Arp2/3 complex activity. B, The activated Arp2/3 complex nucleates the branched actin filament networks for specific formation of the actin cytoskeleton depending on the proper distribution of PI(4,5)P_2_ on the plasma membrane in GH1^GC^. However, failed nucleation of the branched actin networks at the cell cortex as a result of overaccumulation of PI(4,5)P_2_ could disrupt organization of the specific actin cytoskeleton, leading to dysfunctional Golgi apparatus and chloroplasts, subsequently suppressing plant cell elongation in GH1^CB^.

Although SAC phosphatases are essential regulators of PI-signaling networks, little is known about their biochemical and cellular functions in plants. In *Arabidopsis*, AtSac1 has been well characterized as a specific PI(3,5)P_2_ phosphatase localized on the Golgi apparatus, with truncation causing defects in cell morphogenesis and cell wall synthesis (Zhong et al., 2005). Moreover, AtSac7 has been shown to prefer PI4P phosphatase during polarized expansion of root hair cells, thereby regulating the accumulation of PI4P in membrane compartments at the tips of growing root hairs (Thole et al., 2008). Meanwhile, an unknown subgroup of tonoplast-associated enzymes from *Arabidopsis*, SAC2-SAC5, was recently found to be involved in conversion between PI(3,5)P_2_ and PI3P, thereby affecting vacuolar morphology (Novakova et al., 2014). These findings suggest that members of the plant SAC phosphatase family show substrate preference and specific subcellular localization. In contrast, the results of the present study provide evidence that integral ER-localized GH1 phosphatase is required for cell elongation in rice, suggesting a crucial role of SAC phosphatase in crop growth and development. Moreover, it was also observed that the GH1 phosphatase is preferential to bind to and hydrolyze PI4P and PI(4,5)P_2_ (Fig. 2, B-D), suggesting that GH1 specifically regulates metabolism of membrane PI4P and PI(4,5)P_2_, thereby maintaining PI abundance in rice.

Dysfunction of GH1 lead to massive accumulation of PI(4,5)P_2_ and PI4P (Fig. 2, E and F; Supplemental Fig. S3, H and I), while excessive GH1 caused their depletion (Figure S4i,j), suggesting that homeostasis of PIs is pivotal for plant growth and development. In line with this, a previous study revealed that proper balance between PI4P and PI(4,5)P_2_ was necessary for clathrin-dependent endocytosis in the tip of pollen tubes (Zhao et al., 2010), with too much or too little impairing rapid tip growth. This supports the suggestion that flexible turnover of PIs is an important requirement for normal cell morphology. Low abundance and rapid turnover of these lipids provides a sensitive mechanism for fine-tuning protein targeting and function (Boss and Im, 2012). In plants, although activators of the Arp2/3 complex have been somewhat revealed, the determinate counterparts of the Arp2/3 complex remain largely unknown (Frank et al., 2004; Harries et al., 2005; Zhang et al., 2008). Nevertheless, the rice functional WAVE/SCAR protein TUT1 was found to promote actin nucleation and polymerization and contribute to rice development, in which the *tut1* mutants showed defects in the arrangement of actin filaments, and had short roots and degenerated small panicles with reduced plant height (Bai et al., 2015). This study implied that the WAVE/SCAR protein TUT1-activated Arp2/3 complex could control rice plant and panicle development via actin organization in rice, which coincides with the action of *GH1* for rice morphogenesis. In the study, we supposed that overaccumulation of PI(4,5)P_2_ resulted from mutated *GH1* might disrupt Arp2/3 complex-mediated actin polymerization, linking PI metabolism and modulation of the WAVE/SCAR-Arp2/3 complex. These findings at least suggest that the Arp2/3 complex possibly acts as the molecular target of PI(4,5)P_2_-dependent modulation in eukaryotes, providing new insight into the relationship between membrane PI homeostasis and actin cytoskeleton organization. However, despite these findings, the precise molecular mechanisms underlying the action of PM PIs during activation of Arp2/3 complex *in vivo* remains enigmatic. Accordingly, it’s still difficult and time-consuming to identify the mutants of Arp2/3 complex components distinctly in rice, which somewhat limits our advanced understanding of the function of the plants Arp2/3 complex. Nevertheless, future analysis of the membrane PI(4,5)P_2_ and PI4P sensors associated with spatiotemporal colocalization of actin filaments *in vivo* will therefore help determine the relationship between membrane PI homeostasis and actin cytoskeleton organization for cell development.

## Material and Methods

### Plant materials and growth conditions

Parental *indica* variety Guichao-2 (GC) was crossed with *japonica* variety CB to generate F_1_ plants, which were then backcrossed with CB plants to produce BC_1_F_1_ seeds. Repetitive backcrossing and marker-assisted selection generated several plants in which the region around the *GH1* locus was heterozygous, while almost all other regions were homozygous for CB, allowing segregation of populations and fine mapping. A nearly isogenic line (NIL) for *GH1* was subsequently developed from BC_5_F_2_ generations, containing a 50-kb chromosomal region of GC at the *GH1* locus under the CB genetic background. All rice plants were cultivated in experimental fields in Shanghai and Hainan, China, under natural growth conditions.

### Fine mapping of *GH1* and bulked segregation analysis (BSA)

A total of 186 BC_2_F_2_ plants were used to fine map *GH1* at chromosome 2 using grain number per panicle. Subsequently, molecular markers NB-2-80 and NB-2-93 flanking *GH1* were used to assess the recombinants in 6379 BC_3_F_2_ plants. To further determine the location of *GH1*, markers were developed to screen plants showing homozygous recombination in the BC_3_F_3_ progeny. It was then narrowed to a region between the loci defined by markers NB-2-83c and NB-2-83d by identifying the agronomic traits of recombinant plants. Candidate *GH1* genes from GC and CB were then sequenced and compared. The primers used for map-based cloning and the genotyping assays are listed in Table S1. BSA-assisted mapping of *GH1* was performed as described previously (Takagi et al., 2013).

### Transgene construction and plant transformation

The gene editing construct of *GH1* for CRISPR/Cas9 was designed as previously described (Ma et al., 2015). To produce overexpressing transgenic plants, the full-length coding sequence of *GH1* was amplified from GC and cloned into plant binary vector pCAMBIA1306 under control of the CaMV 35S promoter. *Agrobacterium tumefaciens*-mediated transformation of rice with strain EHA105 was then performed as described previously (Hiei et al., 1994). All constructs were confirmed by sequencing. The PCR primers are given in Table S1.

### RNA extraction and qRT-PCR

Total RNA was extracted from different leaves using TRIZOL reagent (Invitrogen). Reverse transcription was performed using ReverTra Ace qPCR RT Master Mix with gDNA Remover (Toyobo) using 500 ng total RNA. Quantitative RT-PCR analysis was carried out using the ABI 7300 Real Time PCR System with Fast Start Universal SYBR Green Master Mix and ROX (Roche), and data were analyzed using the 2^-ΔΔCT^ method. Rice *UBQ5* gene transcripts were used as a reference. The relevant primers for gene amplification are given in Table S1.

### Subcellular localization of GH1

To determine the subcellular localization of GH1, the *GH1* fragment was inserted into the pA7-GFP plasmid and transformed into rice protoplasts. Organelle marker CD3-959-mCherry was used as the ER-marker (Ma et al., 2015). GFP and mCherry fluorescence in rice protoplasts was detected using an LSM 880 confocal laser-scanning microscope (ZEISS). Sequences of the PCR primers used for vector construction are given in Table S1.

### Scanning electron microscopy

For scanning transmission electron microscopy, leaves from 10-day-old rice seedlings grown on 1/2 MS plates were fixed *in situ*. Briefly, the leaves were removed and transferred immediately to vials with fresh primary fixative (1% paraformaldehyde, 2.5% glutaraldehyde, phosphate buffer pH 7.2) followed by 4 h incubation at room temperature. The samples were then rinsed in phosphate buffer and fixed in 1% osmium tetroxide overnight at 4°C before dehydration in an ascending alcohol series. After fixing, the samples were stepwise infiltrated with epoxy resin then embedded in Epon-812 resin before slicing into thin sections using a diamond knife. The samples were then placed on nickel grids, and double-stained with 2% aqueous uranyl acetate and lead citrate for observation under a HITACHI H-7650 transmission electron microscope. For X-ray imaging, the rice culms were fixed in FAA (50% ethanol, 5% glacial acetic acid, 5% formaldehyde) overnight at 4°C then dehydrated in a graded alcohol series before critical point drying. The prepared samples were then examined using a Xradia 520 Versa X-ray imager (ZEISS).

### Protein expression and phosphatase activity assay

The highly efficient Bac-to-Bac baculovirus expression system (Invitrogen) was used for protein expression in Sf9 insect cells. The coding sequences of *GH1* and its mutated version were respectively inserted into the pFastBacHT-C vector. Culturing of the insect cells, and generation of the recombinant bacmid and recombinant baculovirus were performed according to the manufacturer’s handbook. A Mem-PER Plus Membrane Protein Extraction Kit and Ni-NTA Resin (Thermo Scientific) were then used to purify the proteins following the manufacturer’s protocol. To assay specific lipid phosphatase activity *in vitro*, a Malachite Green Assay Kit (Echelon Biosciences) was used to detect the liberated phosphate among different PI substrates according to the manufacturer’s protocol. The primers used for vector construction are listed in Table S1.

### *In vitro* lipid binding assay

For lipid binding assay *in vitro*, the PIP strips (Echelon Biosciences) were blocked in a solution of 5% (w/v) fatty acid-free BSA in Tris-buffered saline plus Tween 20 for 1 h at room temperature, and then the blocked membrane lipid strips were incubated with 0.05 mg/mL fusion proteins for 3 h with gentle agitation at room temperature. The membrane strips were then washed extensively with wash buffer for 3 times. The fusion proteins were detected with HRP-conjugated mouse anti-His monoclonal antibody (ABclonal, AE028, 1:3000 dilution) and visualized by enhanced chemiluminescence.

### *In vivo* measurements of PI4P and PI(4,5)P_2_ levels using ELISA

The PIs were extracted from rice as described previously (Drobak et al., 2000), using 50 mg fresh leaves for each sample. Measurements of PI4P and PI(4,5)P_2_ levels in the different samples were then performed using a PI(4)P and PI(4,5)P_2_ Mass ELISA Assay Kit (Echelon Biosciences) by means of competitive ELISA assay. Based on the assay protocol, the colorimetric signals were read at an absorbance of 450 nm and were inversely proportional to the amount of PI4P and PI(4,5)P_2_ extracted from the cells.

### Staining and visualization of actin filaments

To visualize the actin cytoskeleton in the rice cells, root tips from five-day-old rice seedlings grown on 1/2 MS plates were immersed in PEM buffer (100 mM PIPES, 10 mM EGTA, 5 mM MgSO4, 0.3 M mannitol, pH 6.9) with 2% glycerol. ActinGreen488 Reagent (Thermo Scientific) was added directly to the buffer to stain the actin filaments followed by incubation at room temperature for 35 min. The green fluorescence-labeled actin filaments were then viewed using an LSM 880 confocal laser-scanning microscope (ZEISS). The radar chart based on the MATLAB programming language was used to statistically analyze the direction of the actin filaments (360°).

### *In vitro* actin polymerization assay

To determine the effects of PI4P and PI(4,5)P_2_ on Arp2/3 complex-mediated actin nucleating activity *in vitro*, pyrene-labeled actin assays and monitoring of the branching reaction were performed as described previously (Higgs et al., 1999; Zhang et al., 2010). The purified Arp2/3 protein complex from porcine brain (Cytoskeleton) and GST-tagged VCA domain of human WASP protein (Cytoskeleton) were used to initiate actin nucleation. Monomeric actin (10% pyrene-labeled) was incubated with the Arp2/3 complex and GST-VCA protein in the presence or absence of PI4P and PI(4,5)P_2_ at the indicated concentrations in buffer G (2 mM Tris-HCl pH 8.0, 0.01% NaN_3_, 0.2 mM CaCl_2_, 0.2 mM ATP, 0.2 mM DTT) at room temperature. The time course of actin polymerization was then tracked by monitoring the changes in pyrene fluorescence using a QuantaMaster Luminescence QM 3 PH Fluorometer (Photon Technology International) with excitation set at 365 nm and emission at 407 nm immediately after adding 1 × KMEI buffer (50 mM KCl, 1 mM MgCl_2_, 1 mM EGTA, and 10 mM imidazole-HCl pH 7.0).

### Visualization of actin assembly using TIRFM

Direct visualization of actin assembly was carried out using TIRFM as previously described (Amann and Pollard, 2001; Shi et al., 2013). Briefly, the flow cells were pre-incubated with 25 nM NEM-myosin and equilibrated with 1% bovine serum albumin then washed with fluorescence buffer (10 mM imidazole pH 7.0, 50 mM KCl, 1 mM MgCl_2_, 1 mM EGTA, 50 mM DTT, 0.2 mM ATP, 50 µM CaCl_2_, 15 mM glucose, 20 µg/ml catalase, 100 µg/ml glucose oxidase and 0.5% methylcellulose). Pre-assembled actin filaments (25 nM) in the fluorescence buffer were then injected into the flow cells and allowed to settle. To assess the effect of PIs on actin branching, PI4P and PI(4,5)P_2_ from porcine brain (Avanti) were perfused at various concentrations into the flow cells. The actin filaments were then visualized using an Olympus IX-71 microscope equipped with a ×60, 1.45-numerical aperture Planapo objective (Olympus) by TIRFM illumination. Time-lapse imaging of the actin filaments was captured at 3 sec intervals.

### GST pull-down

The VCA domain of human WASP protein was expressed in a bacterial expression system as a GST-tagged fusion protein (Cytoskeleton). The seven protein subunits of the Arp2/3 protein complex were purified from bovine brain (Cytoskeleton). The GST-VCA fusion protein associated with the glutathione-agarose 4B (GE Healthcare) was incubated with the Arp2/3 complex in pull-down buffer (50 mM Tris-HCl pH 7.5, 150 mM NaCl, 1 mM EDTA, 1 mM DTT, 0.5% Triton X-100 and 5% glycerol) for 2 h at 4°C. Various concentrations of PI(4,5)P_2_ from porcine brain (Avanti) were used for competition. The GST beads were washed and eluted for western blot assays using anti-ArpC2 antibody (Abcam, ab96779, 1:3000 dilution). GST, glutathione S-transferase.

## Acknowledgements

We thank Min Shi (Institute of Plant Physiology and Ecology, SIBS, CAS) for technical support with the transgenic assay. We thank Xiaoyan Gao, Zhiping Zhang, and Jiqin Li (Institute of Plant Physiology and Ecology, SIBS, CAS) for technical support. We thank Chunjie Cao (Carl Zeiss) for X-ray microscopy observation. This work was supported by the grants from the Ministry of Science and Technology of China (2016YFD0100902), National Natural Science Foundation of China (31788103), Chinese Academy of Sciences (QYZDY-SSW-SMC023, XDB27010104), China Postdoctoral Science Foundation (2018M642102), the Shanghai Science and Technology Development (18JC1415000), CAS-Croucher Funding Scheme for Joint Laboratories and National Key Laboratory of Plant Molecular Genetics.

## Author Contributions

H.X.L. conceived and supervised the project, and H.X.L., T.G. and H.C.C. designed the experiments. T.G. and H.C.C. performed most of the experiments. Z.Q.L., M.D., K.C., N.Q.D., J.X.S. W.W.Y., S.J.H. and H.X.L. performed some of the experiments. T.G. and H.X.L. analyzed data and wrote the manuscript.

## Supplemental Data

**Supplemental Figure S1.** Genetic identification of *GH1*. A, Plant architecture of parental varieties GC and CB at the reproductive phase. Scale bar, 10 cm. B, Mature panicles of GC and CB. Scale bar, 5 cm. C, Mature grains from GC and CB. Scale bar, 1 mm. D, The culm of GC and CB plants. Scale bar, 5 cm. E and F, Comparisons between GC and CB for average plant height (n=15 plants) (E), average spikelet number per panicle (n=20 plants) (F), average grain length (n=20 plants) (G), average grain width (n=20 plants) (H). I and L, Comparisons between NIL-*GH1^GC^* and NIL-*GH1^CB^* for average plant height (n=15 plants) (I), average spikelet number per panicle (n=20 plants) (J), average grain length (n=20 plants) (K), average grain width (n=20 plants) (L). Values in (E-L) represent the mean ± SD. \*\**P* < 0.01 compared with the control using the Student’s *t*-test.

**Supplemental Figure S2.** Bulked Segregation Analysis (BSA)-assisted mapping of *GH1*. A, Two subsets of rice with distinct plant height variation, segregated from the same BC_3_F_2_ population, were respectively pooled (>30 plants per pool) and re-sequenced. The calculated SNP-Index is shown. The red and blue lines indicate the pools of low and high plant height, respectively. The green box indicates the region where the SNP-index is divergent between the two pools on chromosome 2. B, Enlarged view of the divergent region in A. The candidate region was between 19.5-22 Mb on chromosome 2.

**Supplemental Figure S3.** Deletion of *GH1* suppresses rice growth and development. A, Plant architecture of GC, *GH1^cas9^-*2, and *GH1^cas9^-*3 at the reproductive phase. Scale bar, 10 cm. B, Mature panicles of GC, *GH1^cas9^-*2, and *GH1^cas9^-*3. Scale bar, 5 cm. C, Mature grains from GC, *GH1^cas9^-*2, and *GH1^cas9^*-3. Scale bar, 1 mm. D-G, Comparisons between GC, *GH1^cas9^-*2, and *GH1^cas9^*-3 for average plant height (n=8 plants) (D), average spikelet number per panicle (n=8 plants) (E), average grain length (n=8 plants) (F), average grain width (n=8 plants) (G). H and I, Comparisons of endogenous levels of PI4P (h) and PI(4,5)P_2_ (i) extracted from rice leaf blades of GC, *GH1^cas9^-*2, and *GH1^cas9^*-3 plants (n=3 biologically independent assays). J-M, Comparisons between GC and *GH1^cas9^-*1 for average plant height (n=15 plants) (J), average spikelet number per panicle (n=15 plants) (K), average grain length (n=15 plants) (l), average grain width (n=15 plants) (M). Values in (D-M) represent the mean ± SD. \*\**P* < 0.01 compared with GC using the Student’s *t*-test.

**Supplemental Figure S4.** Overexpression of *GH1* had a weaker effect on plant and panicle morphogenesis in rice. A, Plant architecture of WYJ (Wu-Yun-Jing 7), *GH1^OE^-*1, and *GH1^OE^-*2 at the reproductive phase. Scale bar, 10 cm. B, Panicles at filling stage of WYJ, *GH1^OE^-*1, and *GH1^OE^-*2. Scale bar, 5 cm. C, Mature grains from WYJ, *GH1^OE^-*1, and *GH1^OE^-*2. Scale bar, 1 mm. D-G, Comparisons among WYJ, *GH1^OE^-*1, and *GH1^OE^-*2 for average plant height (n=10 plants) (D), average spikelet number per panicle (n=10 plants) (E), average grain length (n=10 plants) (F), average grain width (n=10 plants) (G). H, Relative expression levels of *GH1* compared between WYJ and the *GH1^OE^* lines (n=10 plants). The *UBQ5* gene was used as an internal reference to normalize the gene expression data. I and J, Comparisons of endogenous levels of PI4P (i) and PI(4,5)P_2_ (j) extracted from rice leaf blades of WYJ, *GH1^OE^-*1, and *GH1^OE^-*2 plants (n=3 biologically independent assays). Values in (D-J) represent the mean ± SD. \**P* < 0.05; \*\**P* < 0.01 compared with WYJ using the Student’s *t*-test.

**Supplemental Figure S5.** Phylogenetic analysis and amino acid sequence alignment of GH1 homologs from different species. A, Genes showing the highest similarity with *GH1* in *Oryza Sativa*, *Arabidopsis thaliana*, and *Sorghum bicolor* were analyzed using MEGA software. *GH1* was clustered with the type II SAC domain-containing protein family in *Arabidopsis thaliana*. B, Amino acid sequence alignment of the GH1 homologs. The red box indicates the core catalytic motif of GH1, which is absent in the GH1^CB^ allele, while the asterisk indicates the mutated arginine.

**Supplemental Figure S6.** GH1 is a membrane-localized protein. A, Protein sequence analysis using Phobius (http://phobius.sbc.su.se/) predicted that GH1 has two tandem transmembrane domains (grey shade) on the C-terminal. B, Schematic diagram of the transmembrane domains (black block) and core catalytic motif (red block).

**Supplemental Figure S7.** Inhibitory action of PI(4,5)P_2_ towards the Arp2/3 complex is specific, and is released by elevated amounts of VCA. A and B, Spectrofluorimetry assay of actin polymerization using pyrene-labelled actin. a.u., arbitrary units. While 100 μM PI(4,5)P_2_ obviously inhibited Arp2/3 complex activity, approximating the level of actin alone (A), 100 μM PI4P had a much lower inhibiting effect, despite their analogous structures (B). C, In 67 μM fixed PI(4,5)P_2_, VCA released the inhibiting effect of PI(4,5)P_2_ along with elevated amounts, suggesting a competitive relationship between VCA and PI(4,5)P_2_.

**Supplemental Table S1.** Primers used in this study.

**Supplemental Video S1.** Visualization of the Arp2/3 complex-nucleated actin branching networks using TIRFM. The Arp2/3 complex stimulated actin polymerization in conjunction with the indispensable VCA domain of WASP. However, in the presence of 100 µM PI(4,5)P_2_, the actin branches initiated by the Arp2/3 complex were dramatically inhibited, whereas its structural analogue, PI4P, had a much weaker inhibiting effect.

