## Supplementary figures and images for "A SAC phosphoinositide phosphatase controls rice development via hydrolyzing phosphatidylinositol 4-phosphate and phosphatidylinositol 4,5-bisphosphate"

### Supplemental Figure S1

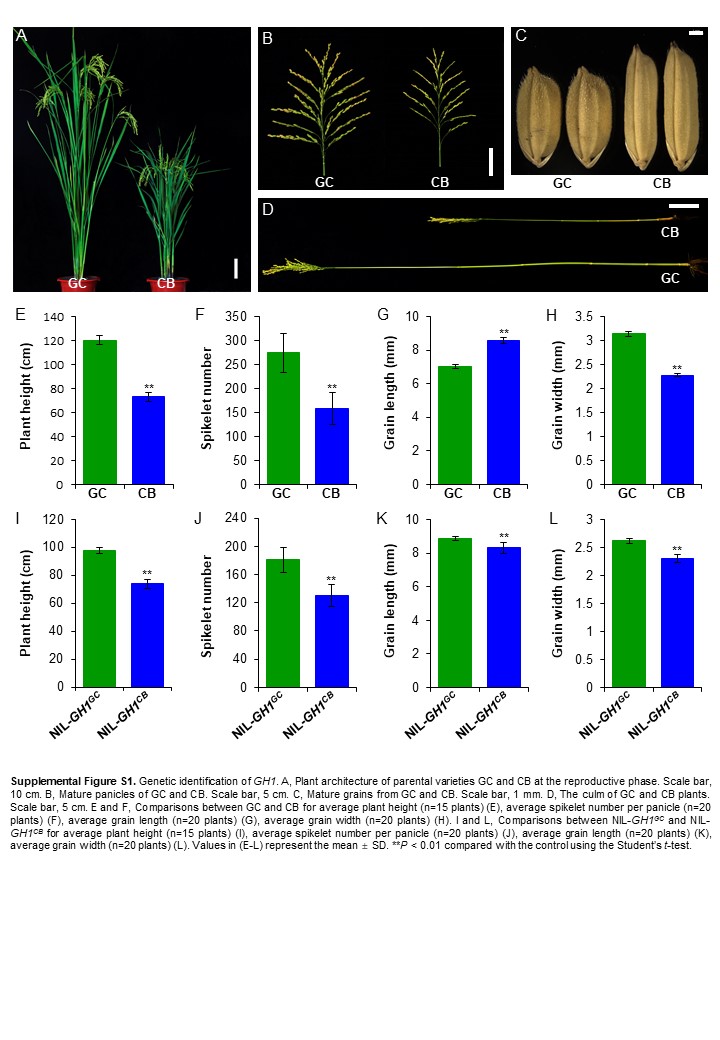

### Supplemental Figure S2

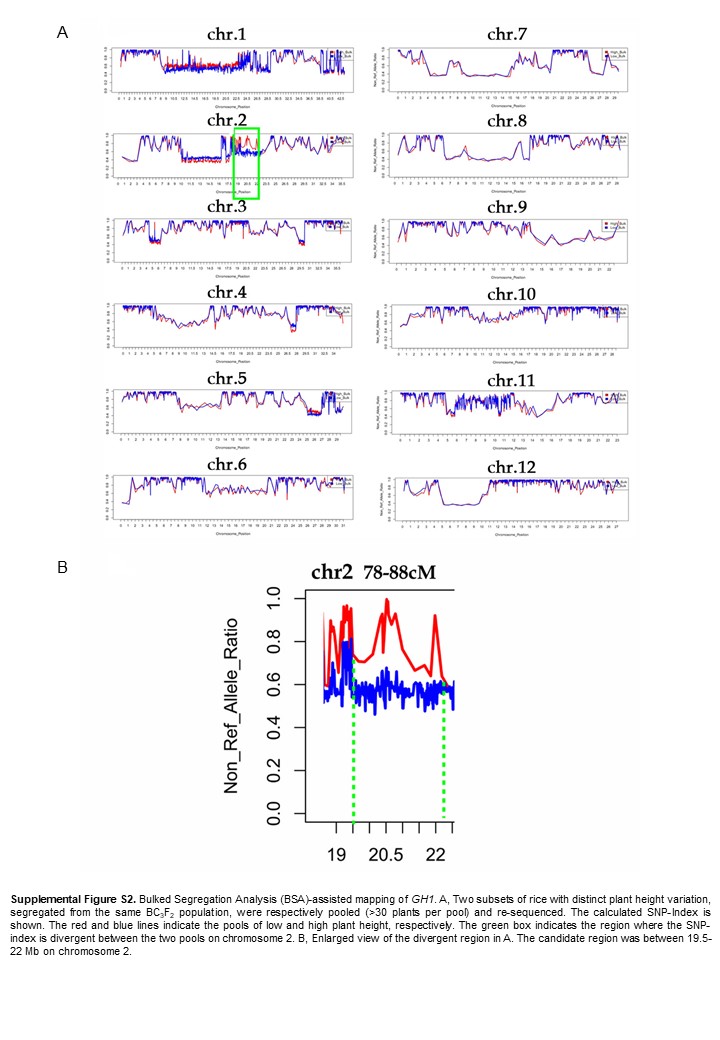

### Supplemental Figure S3

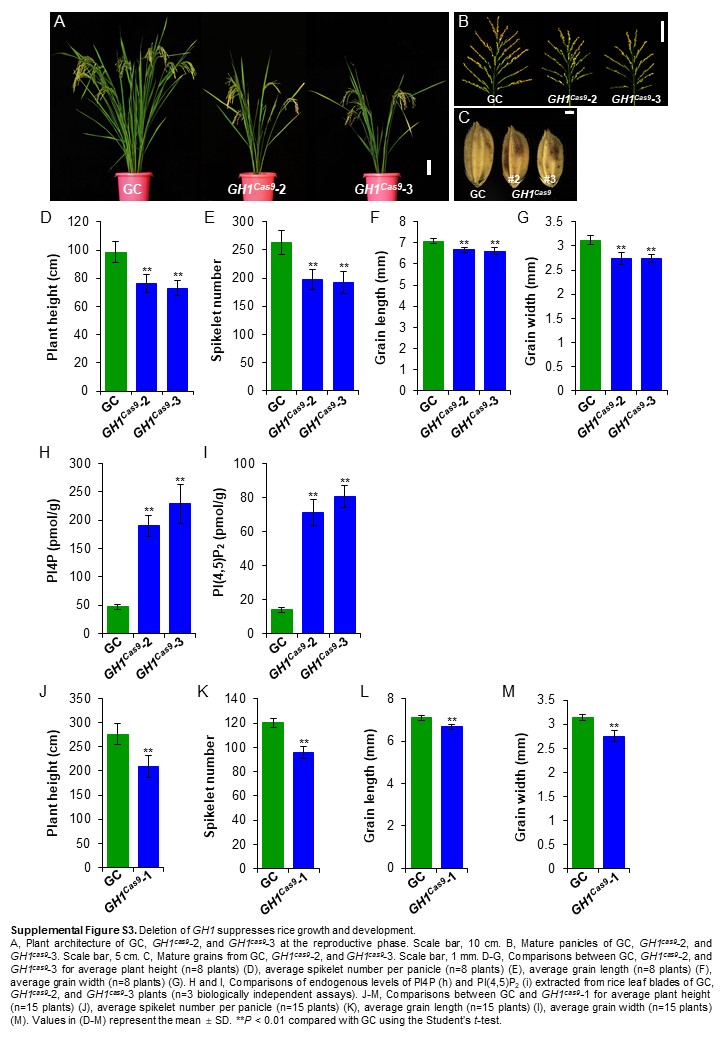

### Supplemental Figure S4

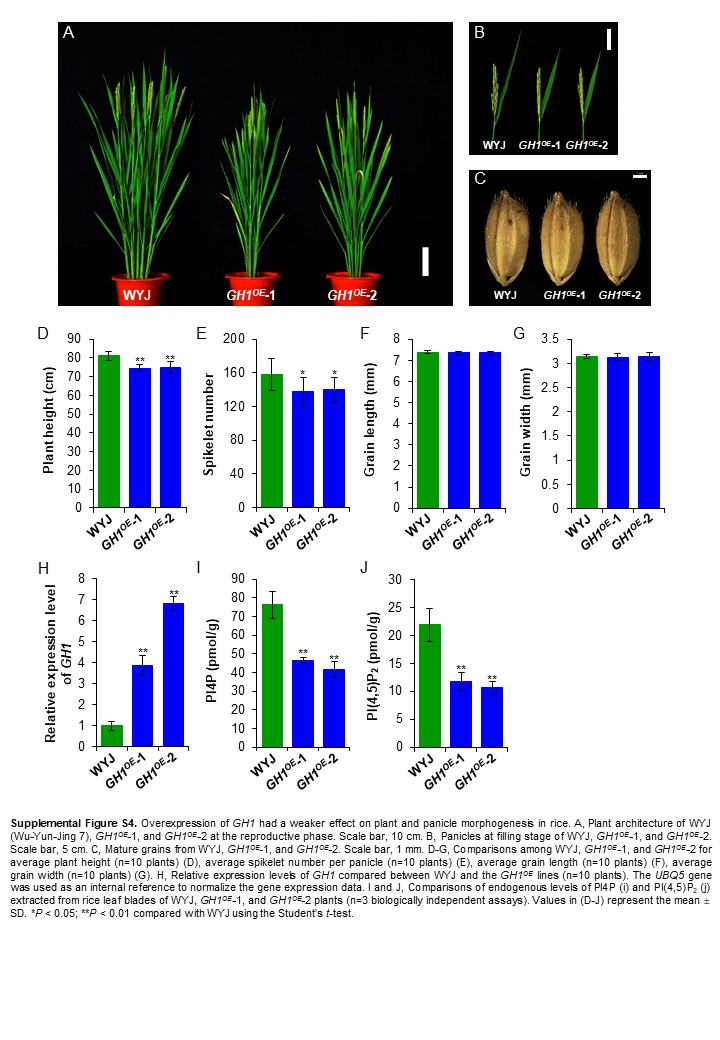

### Supplemental Figure S5

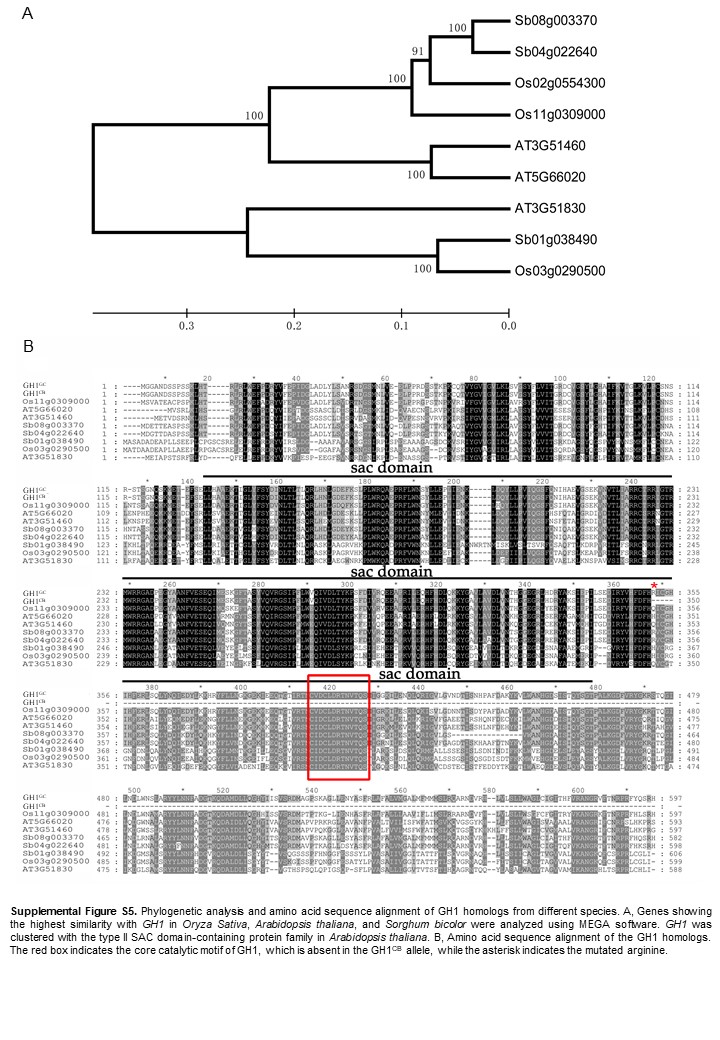

### Supplemental Figure S6

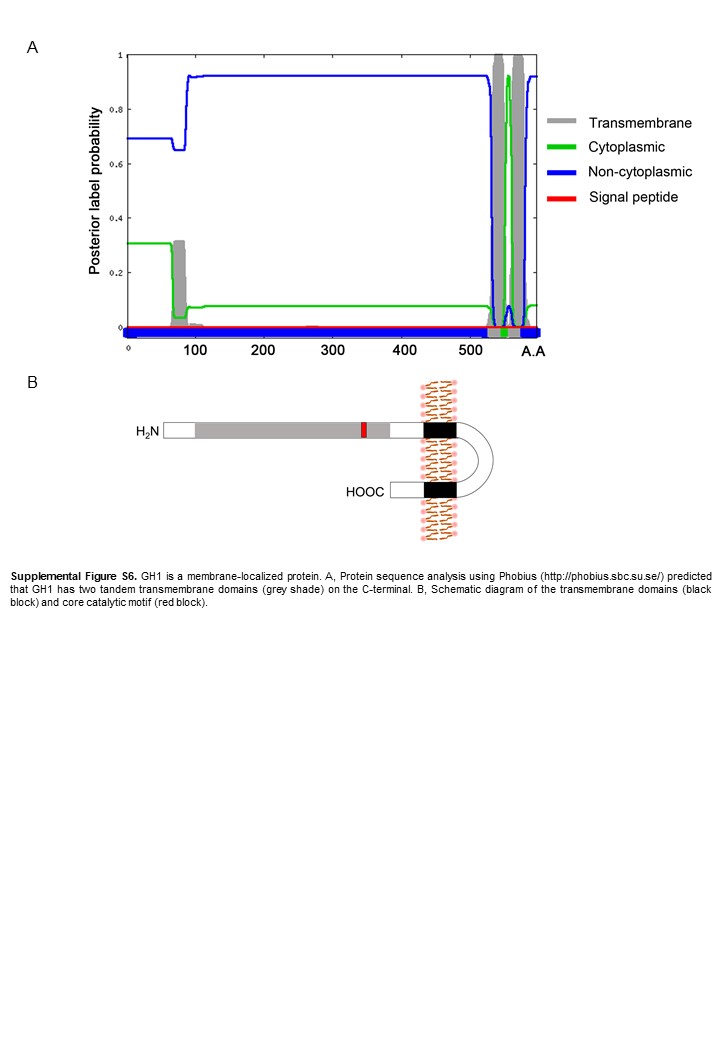

### Supplemental Figure S7

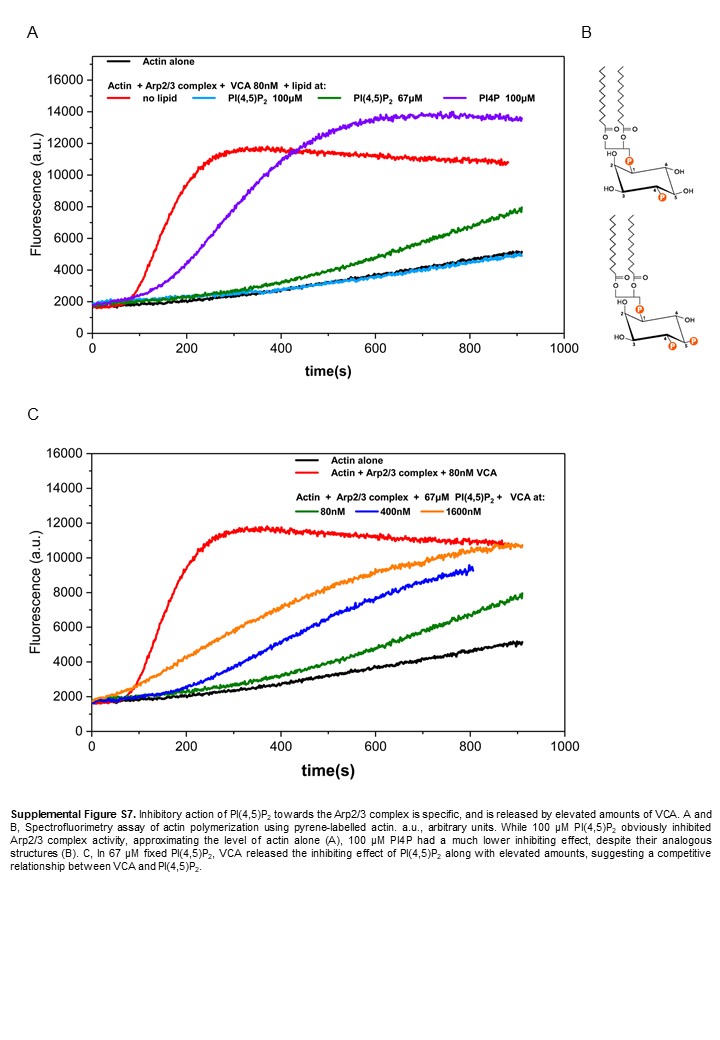
