## Supplemental Table S1 for "A SAC phosphoinositide phosphatase controls rice development via hydrolyzing phosphatidylinositol 4-phosphate and phosphatidylinositol 4,5-bisphosphate"

**Supplemental Table S1. Primers used in this study.**

| Primer Name | Primer Sequence | |
|---|---|---|
| | **Forward (5'-3')** | **Reverse (5'-3')** |
| **Primers for mapping (InDel)** | | |
| RM324 | GATTCCACGTCAGGATCTTCTGG | GCTCACCAGTTGAGATTGAAAGG |
| NB-2-50 | TCTACTGGCATTCTCTCTTTCT | CAAATACCTGTGGTTTTTACCT |
| NB-2-70 | GGGAAAGAGGAGAATTAGAACT | AGTTCTTGATTGTAGAGGCAGT |
| RM341 | CACTCGCGGTTCAAATGCTTACC | TGAACGGCTCCAACGTGAAAGG |
| RM3762 | CCACTAGAATCGGAGAAGTTCACG | TCTACCCTCTATCTCTGGCTCTTCG |
| RM1920 | GCCTGGTAAGTGGTAATGTAATGG | GTGAATTCCTCCTTGGTCTTGG |
| NB-2-80 | CATCCTTTAGTTGACCTTGTCT | CTTTCTGCCGCTTATATCTTAC |
| NB-2-81 | GTCAGACAGAGCGAGTAAAAAG | GGAGAAAATTGCATTATTGTTG |
| NB-2-82 | AATGTTGGCATGGTGTTAAG | GACATCCATGAAACCTGAAA |
| NB-2-83a | CCGTTTCTCTTCTTTTTCCT | TTTCTATCTTGTTCCGTCGT |
| NB-2-83c | AACTAAAGTCACCAGCCAAA | TTGACACAATGCCTGTATTC |
| NB-2-83d | CCGACTTACTAGTGGTGTGTTT | AGGAGTACAACTCTGAACATGC |
| NB-2-85 | ATGGGAATCAAATCAGGACT | CCTCCTTCTATCCACTGCTT |
| NB-2-93 | GAAGCTGTCATGTTCCAGAT | AAGTTGCGCTTAAATACCTG |
| ***GH1* genotyping** | | |
| *GH1*-SNP | TTGGAATCACCTTGTCTTTCG | TTGCTCTTCCCTTTGCCTAC |
| ***GH1* CRISPR/Cas9** | | |
| *GH1*-U3 | GGCAAGTCAGCAAGACCATCAAT | AAACATTGATGGTCTTGCTGACT |
| *GH1*-U6A | GCCGGAGTTCTCAAGCTTTCAGT | AAACACTGAAAGCTTGAGAACTC |
| ***GH1* overexpression** | | |
| p1306-*GH1* | CTAGTCTAGAATGGGTGGGGCAAATGA CTC | ACGCGTCGACATGGCGCGACTGATAA AAAC |
| **Primers for subcellular localization** | | |
| pA7-GH1 | gatactcgagATGGGTGGGGCAAATGACTC | caccatactagtATGGCGCGACTGATAAAAA[illegible] |
| **Primers for protein expression** | | |
| pFastBacHT-GH1 | CGAGCTCACTAGTCGCATGGGTGGGGC[illegible] | cgacaagcttggtacATGGCGCGACTGATA |
| **Primers for qRT-PCR** | | |
| *GH1* | CGTAGCACTTCTGGCAATCA | GTTCTGCCTGTCTCCAAAGC |
| *UBQ5* | ACCACTTCGACCGCCACTACT | ACGCCTAAGCCTGCTGGTT |
